# Leucocyte integrins but not caspases or NLR inflammasome are associated with lipopolysaccharide recognition and response in barramundi (*Lates calcarifer*)

**DOI:** 10.1101/307017

**Authors:** Emmanuelle C. Zoccola, Stuart Kellie, Andrew C. Barnes

## Abstract

The inflammatory response of fish to LPS is subdued, attributed to absence of TLR4, a key pro-inflammatory receptor for LPS in mammals. Nevertheless, LPS is processed in fish in a T-independent manner and is a protective antigen in fish vaccines, yet pathways for processing LPS in fish remain to be elucidated. Here, we report that caspases and NOD-like receptor inflammasomes typically responsible for LPS recognition and processing in mammals lack critical domains or are absent in barramundi (*Lates calcarifer*). However, leucocyte integrins MAC-1 and LFA-1 induce pro-inflammatory cytokine expression poststimulation with LPS. Moreover, MAC-1 and LFA-1 were detected on the surface of neutrophil- and lymphocyte-like cells respectively in the barramundi spleen by immunocytochemistry, and leucocytes displaying MAC-1 or LFA-1 bound to Factor X and ESM-1 respectively. Our results implicate MAC-1 and LFA-1 in immune processing of LPS in barramundi and potentially in antigen processing in fish.

## 1 Introduction

Teleost fish live in permanent contact with relatively high concentrations of bacteria compared with their terrestrial counterparts. A large-scale metagenomics survey by the Tara Oceans project identified that more than 90% of bacteria found in global oceans are gram-negative (Sunagawa et al., 2015). With the exception of the Chloroflexi, all gram-negative bacteria are didermic, comprising an internal cytoplasmic membrane, a thin peptidoglycan layer and an outer membrane comprising lipopolysaccharides (LPS) (Yi and Hackett, 2000). Consequently, fish epithelial and barrier surfaces are continuously exposed to LPS. LPS is typically divided into three structural sections: lipid A, core polysaccharide and repeating O-antigen units (Lüderitz et al., 1982). The LPS lipid A, or endotoxin, is the region of LPS that is recognised by the innate immune system and is highly stimulatory, even at low doses (Miller et al., 2005). LPS recognition in mammals occurs primarily through the toll-like receptor (TLR) 4, which is present on an array of phagocytic immune cells including antigen-presenting cells (APCs) (Poltorak et al., 1998). Briefly, the lipopolysaccharide binding protein (LBP) mediates the interaction between LPS on the bacterial cell surface and the glycoprotein CD14 on phagocytic cells (Wright et al., 1990). CD14 then concentrates LPS to facilitate its binding to the TLR4/myeloid differentiation protein 2 (MD-2), which in turn triggers the inflammatory cascade (Nagai et al., 2002). Lipid A is an essential component of gram-negative bacteria, but it is also highly variable, which can affect its detectability by the immune system (Miller et al., 2005). In fact, there seems to be a correlation between TLR4 recognition of bacterial lipid A and the severity of a disease, with a lipid A poorly recognised by TLR4 more likely to cause severe disease (reviewed in (Miller et al., 2005). In mammals, including humans, lipid A encountered during infections of the bloodstream often causes endotoxic shock, a general inflammatory response which is characterised by fever, hypotension and eventually organ failures that can lead to death (Abbott et al., 2004). Fish, on the other hand, seem to be resistant to endotoxic shock caused by LPS (Iliev et al., 2005b).

Fish show an attenuated regulation of inflammatory cytokines using standard LPS dosages employed in mammalian models, or require approximately 1000 fold higher LPS concentrations to induce a response similar to that observed in mammals (reviewed in (Iliev et al., 2005b). Recently, advances in bioinformatics have established that TLR4 is absent from most of the published genomes and immune transcriptomes from teleost species (Boltana et al., 2011; Iliev et al., 2005b; Zoccola et al., 2017) and, that when present (in *Danio rerio* and *Gobiocypris rarus* for example), the other molecules necessary for recognition of LPS through TLR signalling (LBP, CD14, and MD-2) were absent and/or truncated, non-functional (Iliev et al., 2005b). However, there is evidence of LPS-induced cytokine production in fish (Goetz et al., 2004; Iliev et al., 2005a), which suggest that other molecules are likely involved in LPS recognition and processing in teleosts. In previous work, LPS stimulation induced TNFα transcription through the C-type lectin Mincle in barramundi, but seemed to induce IL-6 transcription through other molecular pathways (Zoccola et al., 2017). Thus, there is potential for alternative LPS receptor families, including inflammatory caspases and leukocyte integrins, to be involved in barramundi leucocyte activation.

Caspases are cysteine proteases that mediate cell death and inflammation in mammals (Sakamaki and Satou, 2009; Takle and Andersen, 2007). Caspases are composed of a CASc domain (comprised of a large p20 and a small p10 subunit), as well as a variable prodomain. Typically, in humans, caspases can be grouped into three sub-categories: cell death initiators, which possess a double death effector domain (DED) motif pro-domain or caspase activation and recruitment domain (CARD) motif pro-domain (caspase-2, −8, −9 and −10); cell death effectors, which only have a short pro-domain (caspase-3, −6 and −7); and inflammatory, which normally possess a CARD motif pro-domain (caspase-1, −4, −5, and - 12). Caspase-4/5, in conjunction with caspase-1/NOD-like receptor (NLR) inflammasomes, are potent contributors to pyroptosis (a type of cell death), mediating the activation of the inflammatory cytokine IL-1β (Baker et al., 2015; Schroder and Tschopp, 2010). Inflammatory caspases have also been shown to stimulate the transcription of nuclear factor-κB (NF-κB), which in turn promotes the transcription of other inflammatory cytokines such as interferons, tumour necrosis factors and interleukins (IL)-6 and −8 (Sollberger et al., 2014).

Leukocyte integrins are transmembrane heterodimeric glycoproteins found on the surface of leukocytes and play a role in several cellular interactions associated with immune functions. The main integrins expressed on leucocytes, β_2_-integrins, are composed of a unique 0 subunit (CD11a, CD11b, CD11c or CD11d), which is non-covalently attached to a common β_2_subunit (CD18) (Arnaout, 2016; Ingalls and Golenbock, 1995). Both CD11b/CD18 (MAC-1) and CD11c/CD18 (p150/95) have been identified as LPS receptors in mammals (Ingalls and Golenbock, 1995; Wright and Jong, 1986). MAC-1 is the most abundant integrin on neutrophils and is also found on natural killer (NK) cells, fibrocytes, B- and T-cells, whereas p150/95 is primarily found on myeloid dendritic cells and macrophages, although it is also found on NK, B- and T-cells (Arnaout, 2016). Moreover, the identification of LPS binding sites on CD18 suggests that β_2_-integrins are able to directly bind and process LPS, translocating NF-κB to the nucleus and inducing inflammatory cytokine release (Ingalls and Golenbock, 1995; Wong et al., 2007; Wright and Jong, 1986). More specifically, MAC-1 has been shown to enable LPS uptake and subsequent inflammatory pathway activation independently of TLR-4 signalling (Scott and Billiar, 2008) and p150/95 has been shown to activate a cellular response after binding to LPS in a CD14-independant manner (Ingalls and Golenbock, 1995).

In the present study, putative barramundi caspases, NLRs and leukocyte integrins were investigated in the barramundi immune transcriptome (Zoccola et al., 2017). Barramundi inflammatory caspases and leukocyte integrins were identified and characterised, providing further insight into the processes underlying LPS recognition in *Lates calcarifer*.

## 2 Methods

### 2.1 Experimental animals and husbandry

Barramundi (*L. calcarifer*) juveniles of approximately 30-50 g were obtained from Australian Native Finfish, Burpengary, Queensland and transported by road to the University of Queensland. Fish were acclimatised for 2 weeks in a recirculating system of eight 84 L cylindrical food-grade plastic tanks with individual aeration, all connected to a 260 L sump equipped with a protein skimmer and a bio-filter. The water temperature and the salinity were maintained at 28 ± 2°C and 15 parts per thousand (ppt) respectively. Water quality was checked regularly for ammonia, nitrite, nitrate and pH, and water exchanges were applied as required. Fish were fed to satiation twice daily with a commercial diet for barramundi (Ridley Aqua Feed). Fish were graded (segregated into different tanks by size) weekly to prevent cannibalism until they reached around 80-90 mm in size, after which they were distributed into their experimental groups.

### 2.2 Bioinformatics analysis

Using the previously generated and annotated barramundi immune transcriptome (Zoccola et al., 2017), several caspases, β_2_-integrins (both 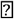 and β subunits) and NLRs were identified by homology in barramundi through local Basic Local Alignment Search Tool (BLAST), using human, murine, crustacean and teleostean caspases sequences (Table S1, S2 and S3 for caspases, NLRs and integrins respectively). Phylogenetic relationships were inferred by maximum likelihood from alignments of barramundi cDNA sequences from the transcriptome, with human sequences listed in Table S1 for caspases and with the cDNA sequences listed in Table S3 for β_2_-integrins. Trees were inferred in MEGA6.06 using ClustalW nucleotide alignment (codon) and supported by 2000 bootstrap replicates. NLR phylogeny was performed using only the proteins’ NACHT domain, identified using the National Center for Biotechnology Information (NCBI) CD-search tool (https://www.ncbi.nlm.nih.gov/Structure/cdd/wrpsb.cgi). Only proteins with a nucleotide-binding and oligomerization domain (NACHT domain) containing the conserved motif G-(4X)-GK-(10X)-W at the N-terminus were used for phylogenetic analysis (Table S2), using the Minimum Evolution method with bootstrap replication (2000) in MEGA6.06, after ClustalW amino-acid alignment (Hughes, 2006). The protein domains were predicted using a combination of SMART (http://smart.embl-heidelberg.de) and CD-search tool on the NCBI website, using each corresponding protein sequence. Functional categorisation for caspases was based on current literature on human and teleostean caspases (Sakamaki and Satou, 2009; Takle and Andersen, 2007).

### 2.3 Adhesion assay

Following the identification of leukocyte integrins, a cell adhesion assay was performed to support the presence of MAC-1 and LFA-1 on barramundi cells. Leukocyte integrins share common ligands but fibronectin, ESM-1 and factor X were identified as specific ligands for the αD, αL and αM subunits respectively (Humphries et al., 2006; Johnson et al., 2009). Collagen was used as a positive control as it has been identified as a ligand for the αL, αM and αX subunits (Arnaout, 2016). Briefly, wells of a high-binding 96 well plate was coated overnight at 4°C with either 100 μL of 20 μg/mL fibronectin, ESM-1, factor X or collagen, or with 100 μL undiluted Poly-L-Lysine solution or 1% BSA as negative controls (all from Sigma). The wells were then washed three times in 1M phosphate buffer saline (PBS) before blocking for 1 h at 37°C with 1% bovine serum albumin (BSA) to prevent non-specific binding to the plastic. Half the wells containing ESM-1 and factor X were simultaneously incubated with antibodies specific for Integrin αM (ITGAM) and integrin αL (ITGAL) subunits respectively, as supplementary controls. After the wells were washed thrice in PBS, cells were isolated from spleen by passing the organ through a 100 μm mesh and subsequently washed in RPMI by centrifugation (300 × *g*, room temperature (RT), 5 min). The cells were then resuspended in RPMI at 10^6^ cells/mL and seeded at 100 μL/well. After 1 h incubation at 28°C, the wells were inverted to remove the media and non-adherent cells before being washed twice using 300 μL of ice-cold 1M PBS containing 1 mM CaCl_2_ and 1 mM MgCl_2_. The cells were then fixed and permeabilised using ice-cold methanol for 10 min at RT. After three washes in 1M PBS, the wells were stained at RT for 10 min using a crystal violet solution (0.5%w/v crystal violet in 20% ethanol). After three washes by immersion in deionised water, the crystal violet retained was dissolved using 150 μL of 100% methanol for 15 min at RT and quantified by absorbance at 590 nm with a Fluostar Optima plate reader (BMG Labtech, Melbourne, Australia).

### 2.4 Immunohistochemistry and fluorescent microscopy

Glass coverslips were sterilised using ethanol and a flame, and were placed in each well of a 24-well plate. The coverslips were then coated overnight at 4°C with either 20 μg/mL fibronectin, ESM-1, factor X or collagen, or with undiluted Poly-L-Lysine or 1% BSA as negative controls. Spleens were homogenised into cell suspensions as previously described (Zoccola et al., 2017) and leucocytes were isolated on a Percoll gradient as adapted and modified from Tumbol et al. (2009). Briefly, the cell suspension was layered over a 34%-51% discontinuous Percoll gradient and centrifuged for 30 min at RT (800 × *g*, acceleration 6, brake 0, Eppendorf 5810R). The buffy layer between the two Percoll densities was collected and washed twice in PBS by centrifugation (400 × *g*, acceleration 9, brake 9, RT, Eppendorf 5810R), before being resuspended in L-15 with 5% heat inactivated barramundi serum and incubated in a 6-well plate overnight at 28°C. The next day, the 24-well plate containing the cover slips was washed thrice in PBS and blocked with 1% BSA at 37°C for 1 hour. Cells incubated overnight were washed in PBS to remove serum and concentration was adjusted to 2×10^6^ cells/mL and the cells were plated into the 24-well plate. The plate was incubated for 4 hours at 28°C before the coverslips were fixed in PFA overnight at 4°C.

The coverslips were then washed three times in PBS and permeabilised in Triton X for 3 min at RT, before being washed again thrice in PBS. Coverslips were then blocked in 1% BSA for 1 hour, and following a further wash in PBS, they were incubated with either rabbit-anti-ITGAM or rabbit-anti-ITGAL primary antibodies at 1:100 or with PBS as a negative control for 4 hours at RT. After a further three PBS washes, coverslips were incubated with goat-anti-rabbit IgG conjugated with Alexa Fluor 488 (1:500) in the dark at RT. After 1 h, DAPI was added at 5 μg/mL and the coverslips were incubated for a further 30 min for a total incubation time of 1 h 30 min.

Coverslips were viewed with an Olympus BX41 epifluorescent microscope. Images were captured with an Olympus DP26/U-CMAD3 camera and optimised with the imaging software CellSens (Olympus Optical Co. Ltd, Japan).

### 2.5 Inflammatory cytokine regulation assessed by qRT-PCR

TNFα, IFN-α, NF-κB, IL-1β and IL-6 were chosen for assessment by qRT-PCR due to their inducible nature by LPS through processing by β-integrins (Ingalls and Golenbock, 1995; Wong et al., 2007; Wright and Jong, 1986). Spleens were sampled aseptically from healthy juvenile barramundi and a cell suspension was obtained as described above. Cells were incubated with LPS (*E. coli* 0111:B4) at 0.05 μg/mL for 1 h at 28°C. Blank controls consisted of cells alone, antibody controls consisted of cells incubated with 1 μg/mL of mouse anti-ITGAM or mouse anti-ITGAL blocking antibody for 1 h at 28°C followed by a 1 h incubation with 0.05 μg/mL of LPS at 28°C. Positive controls consisted of cells incubated with 1 μg/mL of phorbol 12-myristate 13-acetate (PMA) for 1h at 28°C. Cells from all treatment groups were then harvested in RNAlater. For each sample, RNA was extracted using the RNeasy kit (Qiagen) according to the manufacturer’s instructions. Contaminating genomic DNA was removed by on-column digestion with the RNAse-Free DNAse set (Qiagen) and the resulting RNA was converted to cDNA from a total of 12 ng of starting RNA per sample using the QuantiTect Reverse Transcription kit (Qiagen). Subsequently, the cDNA was used to assess the relative expression of TNFα, IFN-α, NF-κB, IL-1β and IL-6 by qRT-PCR, using elongation-factor-1-α and Syk as normalisers, as they present the most stable expression levels across treatments and presented similar efficiencies (within 10% of target genes and of each other), on a ABI-ViiA7 cycler (Applied Biosystems). The primers were designed to span across exons when possible to eliminate gDNA amplification (Table 1). After primary optimisation, only TNFα, NF-κB and IL-1β expression levels were assessed, as the primers for IFN-α and IL-6 could not be optimised to the template.

### 2.6 Statistical analysis

Prior to qRT-PCR analysis, the stability of the internal control genes was assessed and the relative expression for each gene was computed using the Relative Expression Software Tool (REST) (Pfaffl,2001; Pfaffl et al., 2002; Schmittgen and Livak, 2008). Data from the adhesion assay were analysed using multiple T-tests in GraphPad Prism, with statistical significance determined using the Holm-Sidak method, with alpha = 0.05. Each row was analysed individually, without assuming a consistent SD. Data from qRT-PCR were analysed through REST.

### 2.7 Ethics statement

All animal work was conducted in accordance with Animal Care and Protection Act and the NHMRC Code of Practice. Work was conducted under the University of Queensland Animal Ethics Committee Approval No. NEWMA/078/15 “Understanding the early onset of adaptive immunity in fish.”

## 3 Results

Putative barramundi caspases were identified by homology and in most cases grouped clearly with their human counterpart (Figure 1). However, while some barramundi caspases had several isoforms (caspase 3 and caspase 2) the inflammatory caspase 4 and the differentiation caspase 14 were not identified in barramundi. Moreover, although identified by homology in barramundi, caspase 5 and caspase 9 were lacking a CARD pro-domain, thus differing from their human counterparts (Figure 1).

**Figure 1.**
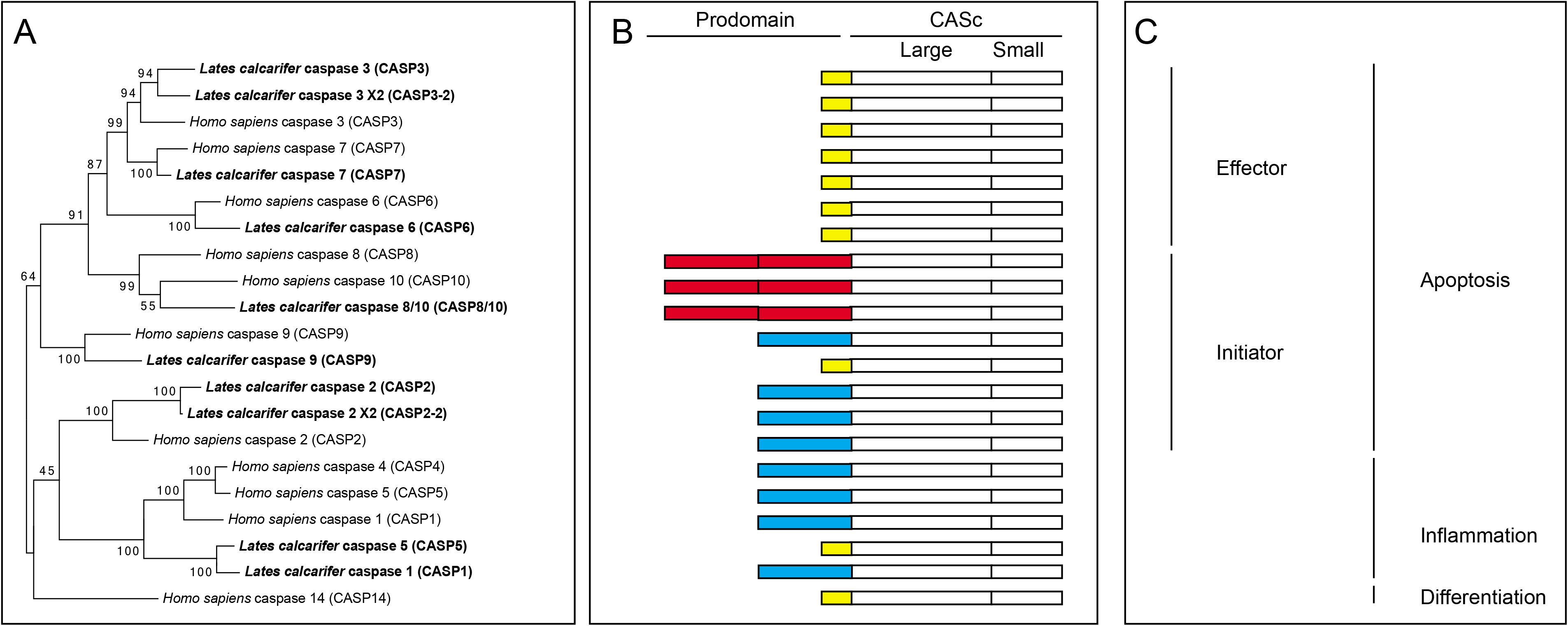
Phylogenetic relationship between human and barramundi (in bold) caspases (A), linked with domain organisation of each corresponding protein (B) and functional categorisation based on literature (C). Yellow boxes correspond to a short pro-domain; Red boxes correspond to a DED (death-effector domain) protein domain; Blue boxes correspond to a CARD (caspase activation and recruitment domain) protein domain.

Putative barramundi NLRs were identified by homology and grouped with their human and murine counterparts (Figure 2). Out of the three distinct NLR sub-families (nucleotide-binding oligomerization domain (NOD), NLR family, pyrin domain containing (NLRP) and Ice protease-activating factor (IPAF)) (Schroder and Tschopp, 2010), only sequences coding for protein from the NOD sub-family were identified in barramundi, with members from the NLRP and IPAF families lacking. Although most protein domains were conserved between human, mouse and barramundi NLRs, some mouse and barramundi proteins were lacking either the pyrin domain (PYD) or caspase activation and recruitment domain (CARD) at the N-terminus or were lacking leucine-rich repeat (LRR) domains at the C-terminus (Figure 2).

**Figure 2.**
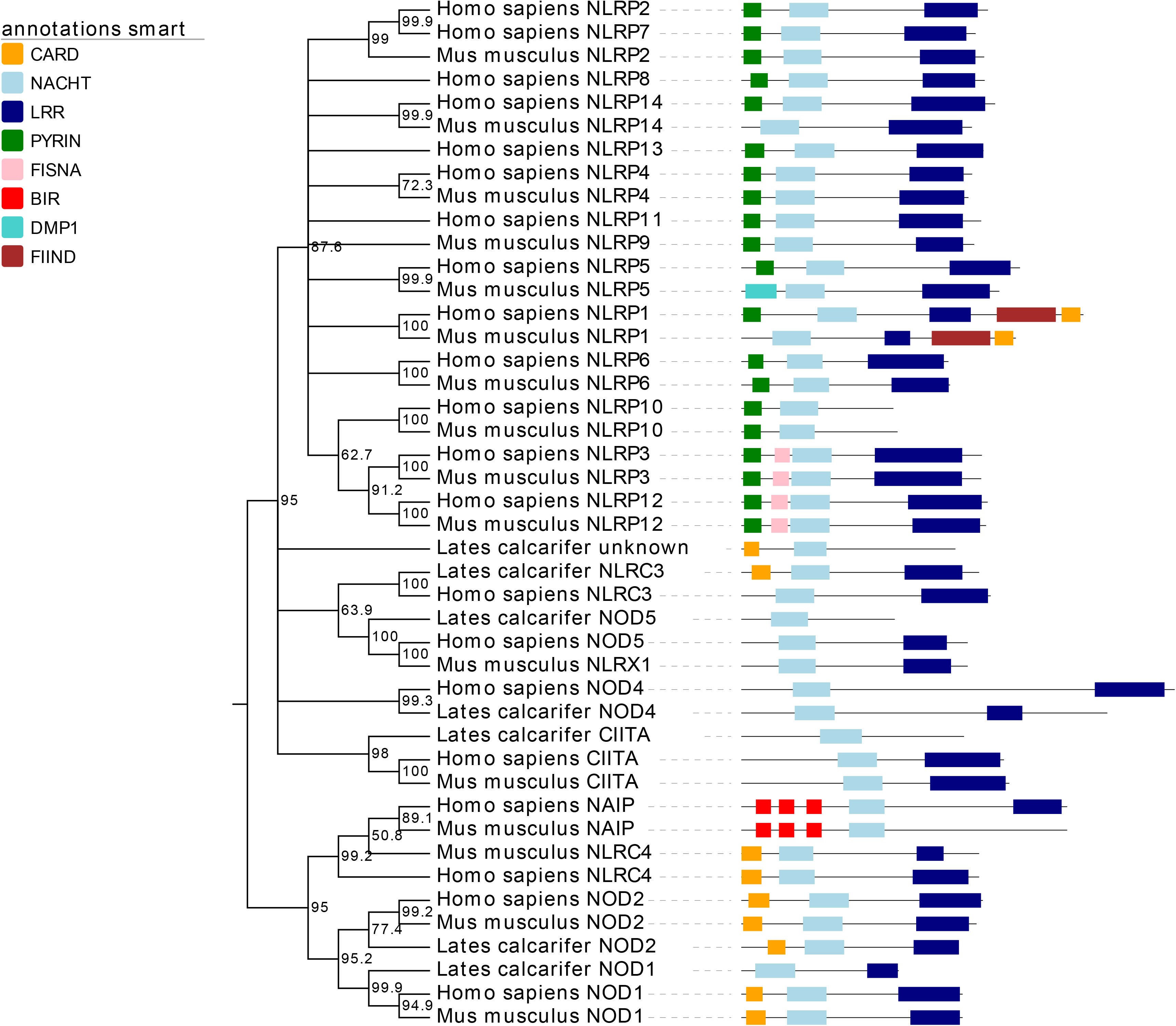
Phylogenetic relationship between human, murine and barramundi (in bold) NLRs based on their NACHT domain. Protein domain organisation is depicted on the right of the phylogenetic tree. CARD: caspase activation and recruitment domain; NACHT: nucleotide-binding and oligomerization domain; LRR: leucine-rich repeat; PYR: pyrin domain; FISNA: Fish-specific NACHT associated domain ; BIR: baculoviral inhibition of apoptosis protein repeat domain; DMP1: Dentin matrix protein 1 ; FIIND: domain with function to find.

Putative barramundi α- and β2-integrin subunits were identified by homology with their human, murine and teleost counterparts (Figure 3). Out of the four possible α subunits forming leukocyte integrins (D, M, L and X), only two were identified in barramundi: M (forming MAC-1) and L (forming LFA-1). Both include a Von Willebrand factor type A, which is required for metal ion ligand binding, and several β-propellor repeats. Moreover, the putative barramundi integrin αM identified was missing a transmembrane domain, which suggests that integrin αM may be secreted rather than membrane bound in barramundi.

**Figure 3.**
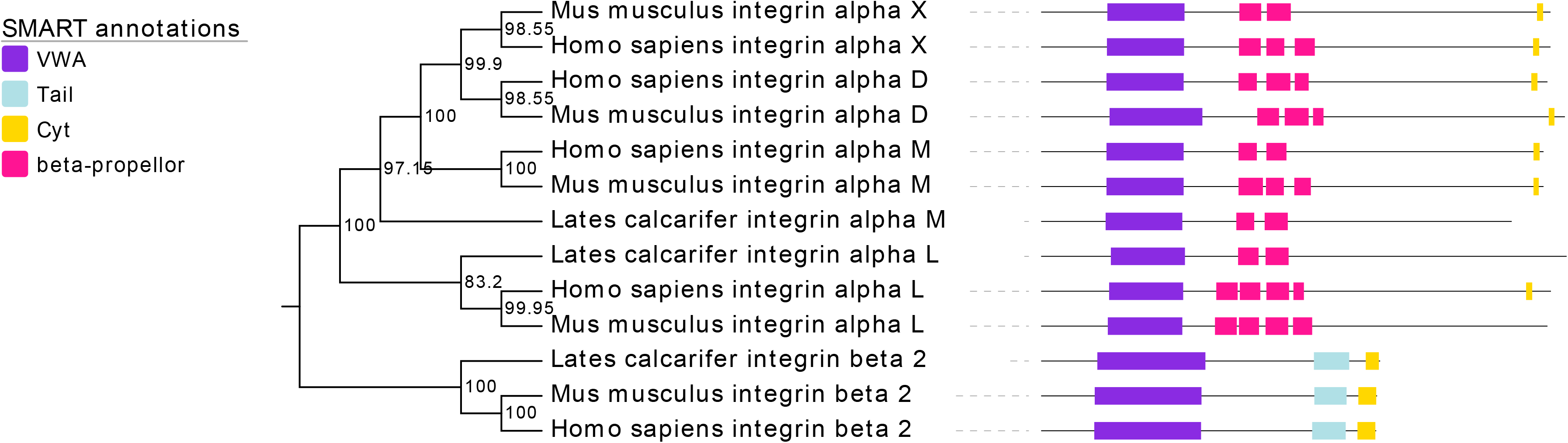
Phylogenetic relationship between human, murine, teleost and barramundi (bold) α- and *β*2-integrin subunits. Protein domain organisation is depicted on the right of the phylogenetic tree. VWA: Von Willebrand factor type A; Tail: tail domain; Cyt: cytoplasmic domain.

When incubated with substrates specific to the D, M and L α-sub-units, barramundi spleen leucocytes did not bind significantly to any of the negative controls or to fibronectin (specific substrate for integrin αD subunit) but did bind significantly to factor X and ESM-1 (specific substrates for integrin αM and L respectively) (Figure 4). Interestingly, when incubated with anti-ITGAM antibody, cells from barramundi spleen were more adherent to the integrin αM substrate factor X (Figure 4). Similarly, cells incubated with anti-ITGAL antibody were more adherent to integrin αL substrate ESM-1 (Figure 4).

**Figure 4.**
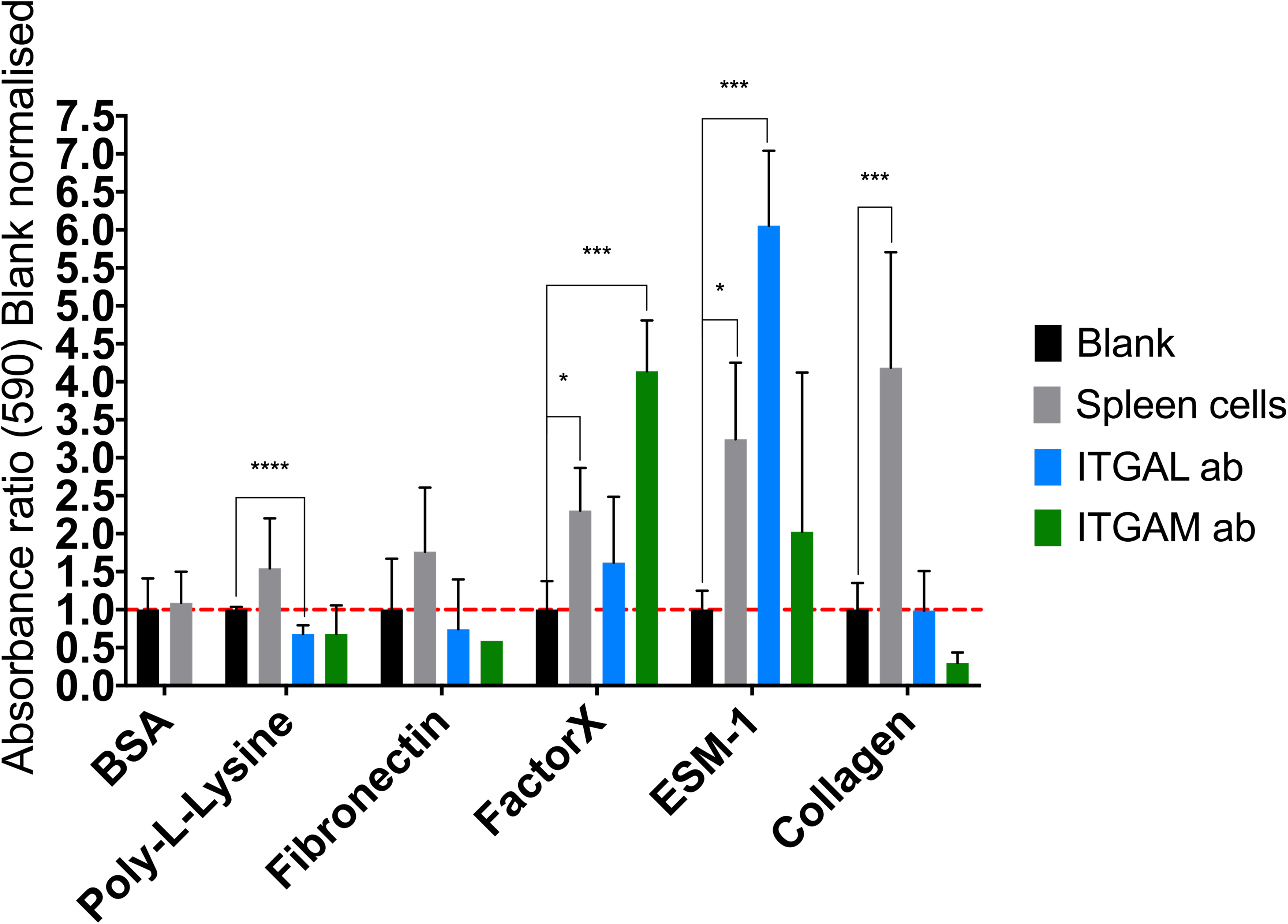
Adherence of barramundi spleen cells to different substrates. Significant differences between blank incubated and cells incubated wells are represented with * p < 0.05, ** p < 0.01, *** p < 0.005 and **** p < 0.001.

The numbers of barramundi spleen leucocytes bound to the positive control, collagen (specific substrate for integrin αM, L and X subunits), was also significantly higher than BSA-coated control.

When the adherent cells were observed by microscopy, the cells binding to Factor X were larger and more granular, resembling granulocytes whereas cells binding to ESM-1 were rounded and slightly smaller, resembling lymphocytes (Figure 5).

**Figure 5.**
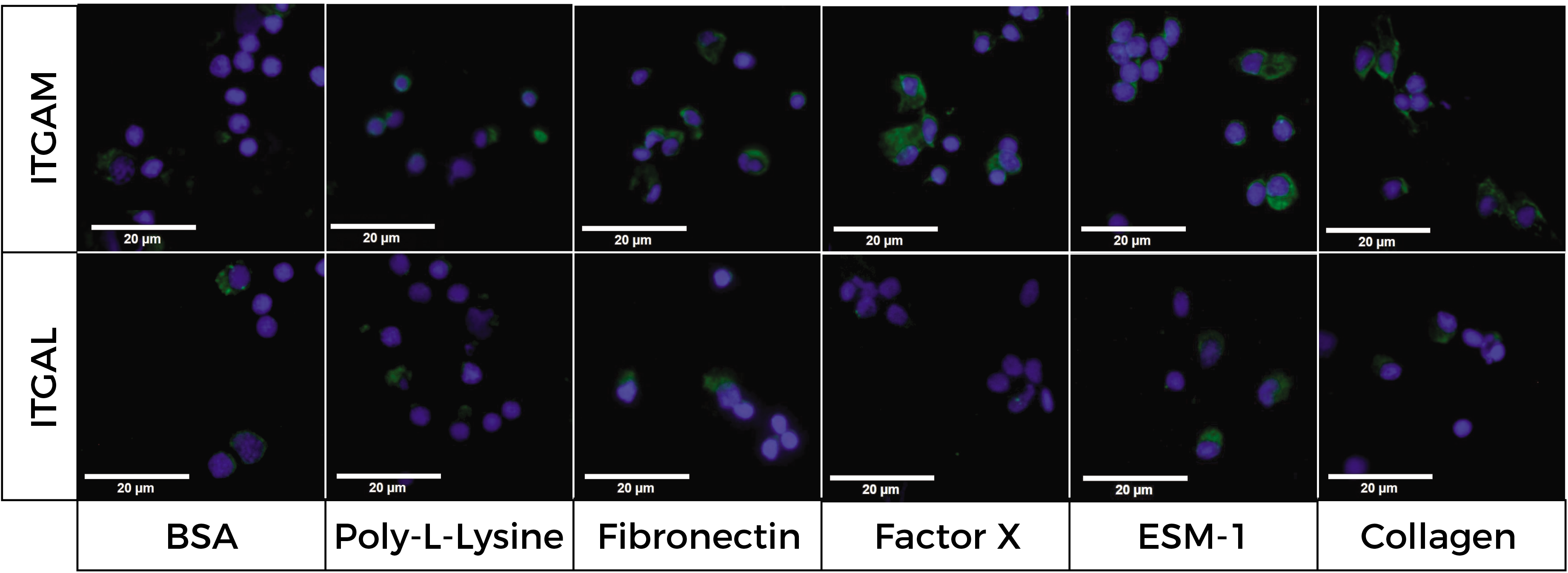
Micrographs of the cells adhering to each substrate, with DAPI stain (nucleic acid) shown in blue and ITGAM or ITGAL specific antibodies stain shown in green.

To determine whether integrins might be involved in LPS recognition and response, a stimulation assay was conducted and the expression of a cohort of cytokines determined by qPCR. PMA (pro-inflammatory positive control) stimulation downregulated the expression of IL-1β but significantly upregulated the expression of TNFα and NF-κB (Figure 6). LPS stimulation, on the other hand, upregulated the expression of IL-1β but did not impact the expression of TNFα or NF-κB (Figure 6). In general pre-incubation with antibody did not significantly affect stimulation by PMA or LPS, with the exception of IL-1β, which was significantly down regulated by PMA stimulation, but this was blocked by incubation with anti-ITGAL antibody (Figure 6A).

**Figure 6.**
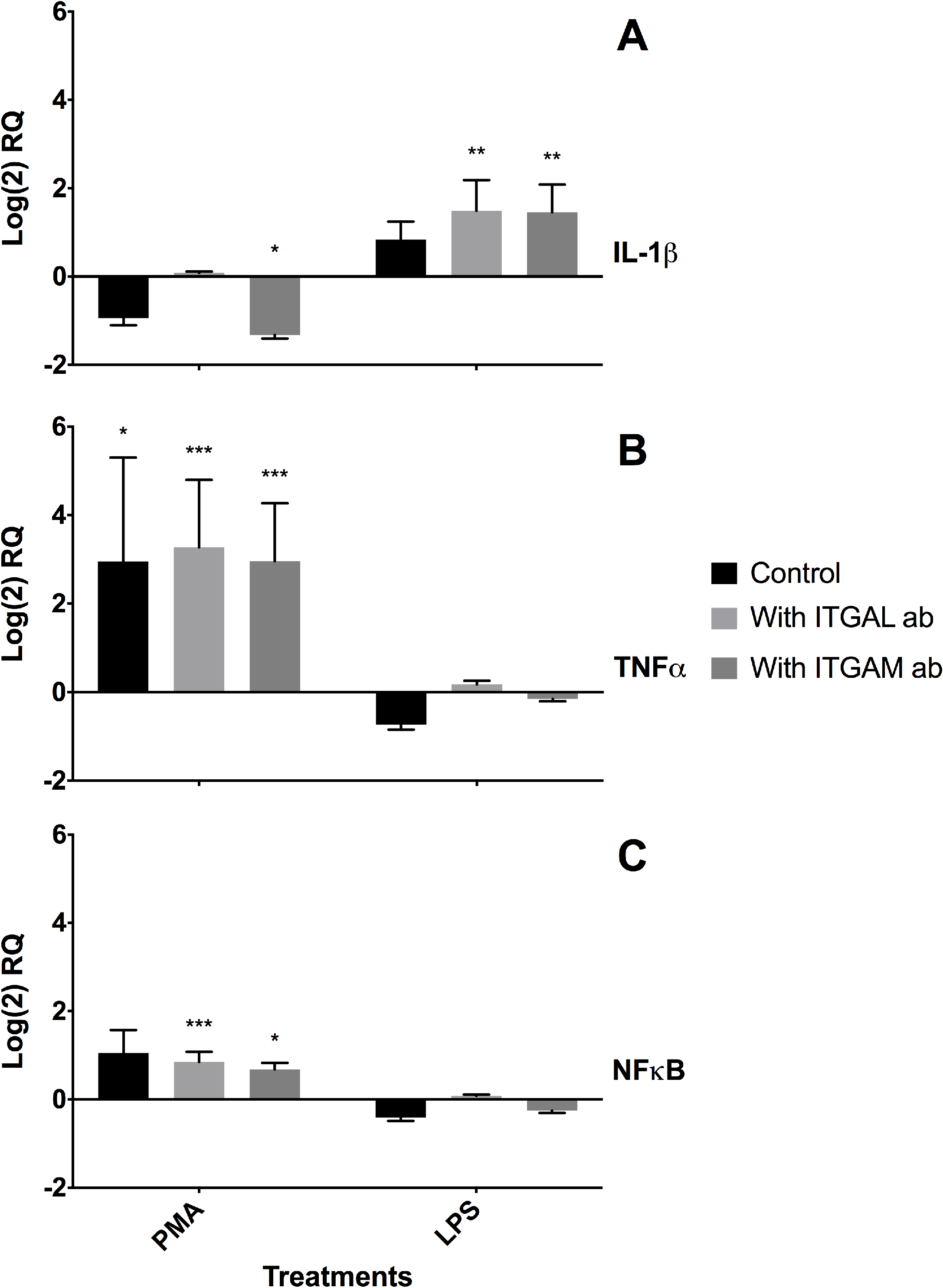
Relative expression of IL-1β (A), TNFα (B) and NF-κB (C) in barramundi splenocytes after stimulation with PMA or LPS with or without pre-incubation with anti-ITGAL or anti-ITGAM antibody. Significant differences between control unstimulated cells gene expression and stimulated cells are represented with * p < 0.05, ** p < 0.01 and *** p < 0.005.

## 4 Discussion

Inflammatory caspases have recently been shown to recognise and process intracellular LPS in humans (caspase-4 and −5) and in mice (caspase-11), binding to LPS through their CARD domain (Baker et al., 2015; Shi et al., 2014). In barramundi, the inflammatory caspases −5 and −1 were identified, and grouped tightly with their human counterparts. However, the CARD domain from barramundi caspase-5 was lacking, suggesting a potential loss of function as CARD is necessary for LPS recognition and CARD oligomerisation is also necessary for caspase activation (Reis et al., 2012). Another way for caspases to recognise PAMPs is through the coupling of caspase-1 with other specific pattern recognition molecules (mostly NLRs) through their CARD domain, forming functional inflammasomes (Petrilli et al., 2007; Sollberger et al., 2014). As barramundi caspase-1 displayed a CARD pro-domain, inflammasomes were plausible actors in LPS recognition so NLRs and other inflammasome sensor proteins were subsequently investigated in barramundi. To date, there are six characterised NLR-inflammasomes, formed using six different NLRs: NLRP1, NLRP3, NLRP6, NLRP7, NLRP12 and NLRC4/IPAF (Latz et al., 2013; Schroder and Tschopp, 2010; Sollberger et al., 2014). Other molecules such as RIG-I, AIM2 and IFI16 have also been described as inflammasome sensor proteins but recognise nucleic acid, rather than carbohydrate-based ligands such as LPS (Latz et al., 2013). None of the NLRs involved in inflammasomes were identified in *L. calcarifer* in the current study, suggesting that inflammasomes in barramundi are not involved in detecting LPS thereby implicating other pathways in LPS recognition in these fish. As inflammasomes typically trigger an inflammatory response, the lack of NLR inflammasomes in barramundi correlates with the weak inflammatory response observed in barramundi and other fish post-stimulation with LPS (Zoccola et al., 2015).

Leukocyte integrins are found at the surface of white blood cells and are involved in pathogen recognition through conformational changes after activation by ligands (Harris et al., 2000). Out of the four possible genes coding for leukocyte integrin α-subunits, two were identified in barramundi: the subunit αM (CD11b) which forms MAC-1 and the subunit αL (CD11a) which forms LFA-1. Cells incubated with Factor X, a specific ligand for the integrin αM subunit (Humphries et al., 2006), bound significantly more to Factor X than to negative control substrates. This suggests that barramundi leucocytes express a protein that is conformationally similar to the αM subunit of MAC-1, hence able to bind to Factor X. The same was true for cells incubated with ESM-1, a specific ligand for the integrin αL subunit (Humphries et al., 2006), suggesting that barramundi leucocytes also produce a protein with binding sites similar to the αL subunit of LFA-1. Incubation of spleen cells with anti-ITGAL or anti-ITGAM antibody increased adhesion of barramundi spleen cells to the corresponding substrates. The antibodies may activate β2-integrins by configurational change, increasing the molecules’ binding strength as previously reviewed (Gahmberg et al., 1998). Indeed, integrin αMβ2 (MAC-1) and αLβ2 (LFA-1) can assume two conformations, open (active) and closed (inactive), which differentially recognise ligands (Abram and Lowell, 2009; Harris et al., 2000). Changes in configuration are crucial to integrin function and can also influence avidity and affinity of the leucocyte integrin (Harris et al., 2000; Hogg et al., 1993), impacting adhesion to ligands, as detected in the adhesion assay reported here (Figure 4). Currently, two binding sites for LPS have been identified on the β2 chain of leucocyte integrins (Wong et al., 2007). Moreover, integrin αMβ2 has been shown to enable LPS recognition and activation independently of TLR4 (Scott and Billiar, 2008), and processes depending on MAC-1 activation have been shown to induce the inflammatory cytokine IL-1β’s expression (Fuhlbrigge et al., 1987; Scott and Billiar, 2008). When observed by fluorescence microscopy, most spleen cells that bound to Factor X expressed integrin αM (ITGAM) and resembled granulocytes, being larger and more granular than cells not expressing ITGAM, similarly to previous observations in peritoneal cells (Arnaout, 1990; Ghosn et al., 2008). Almost all cells that bound to ESM-1 expressed integrin αL (ITGAL) and were more rounded, resembling B- or T-lymphocytes (Arnaout, 2016). Considering that MAC-1 has recently been identified as a LPS receptor in mammals (Scott and Billiar, 2008), and that barramundi leucocytes express MAC-1, it is possible that LPS in barramundi is processed through MAC-1. The expression of inflammatory cytokines following stimulation with LPS was thus investigated by qRT-PCR in barramundi spleen cells. Although the expression of IL-1β was higher in cells stimulated by LPS, it was not statistically significant within the power of this experiment, and LPS did not affect TNFα or NF-κB expression. Prior incubation with anti-ITGAL and -ITGAM antibodies, however, activated integrins on barramundi splenocytes evidenced by significantly increased expression of IL-1β post-stimulation with LPS (Figure 6A). Stimulation with PMA, on the other hand, bypasses cellular receptors and activates protein kinase C directly (Weiss et al., 1984). PMA resulted in downregulation of IL-1β and in consistently higher expression of the pro-inflammatory cytokine TNFα, as well as higher expression of NF-κB (Figure 6B-C) in barramundi splenocytes. Most likely, PMA stimulation thus activates inside-out integrin signalling, through receptors other than the integrin itself, resulting in integrin activation and increased binding strength (Abram and Lowell, 2009). Additionally, stimulation of a mixed leukocyte population with PMA upregulates the expression of TNFα (Schutte et al., 2009), which has been shown to activate NF-κB expression through integrin signalling (Kettritz et al., 2004). LPS, on the other hand, activates outside-in signalling, binding directly to the integrin receptor and activating caspase-1, resulting in IL-1β upregulation but not impacting the expression of TNFα (Walzog et al., 1999). However, before integrins can act as receptors, they need to be activated, which was likely effected by incubation with ITGAL and ITGAM antibodies prior stimulation, supporting the difference in IL-1β regulation by LPS between pre-stimulated and control cells (Figure 6A).

The C-type lectin Mincle was previously identified as a receptor for LPS in barramundi, but other unidentified receptors were also implicated in recognition of bacterial polysaccharides in perciform fish (Zoccola et al., 2017). In the current study, we show that that leucocyte integrins αMβ2 (MAC-1) and αLβ2 (LFA-1) are present on barramundi (*L. caỉcarifer*) leucocytes and play a role in LPS recognition and processing. The absence of some caspase CARD domains and NLRP is also indicative the inflammasome formation may not be possible in this species. Further investigation is therefore warranted amongst the teleostei to determine whether inflammasome formation arose later during vertebrate evolution.

## References

Abbott, J.D., Ball, G., Boumpas, D., Bridges, S.L., Chatham, W., Curtis, J., Daniel, C., Hughes, L.B., Kao, A.H., Langford, C, Lovell, D., Manzi, S., Müller-Ladner, U., Patel, H.C., Roubey, R.A.S., Saag, K., Sabatine, J.M., Shanahan, J., Simms, R., Smith, E., Sundy, J., Szalai, A.J., Wimmer, T., 2004. Endotoxic shock, in: Moreland, L.W. (Ed.), Rheumatology and Immunology Therapy. Springer Berlin Heidelberg, Berlin, Heidelberg, pp. 302–303.

Abram, C.L., Lowell, C.A., 2009. The ins and outs of leukocyte integrin signaling. Annu Rev Immunol 27, 339–362.

Arnaout, M.A., 1990. Structure and function of the leukocyte adhesion molecules CD11/CD18. Blood 75, 1037–1050.

Arnaout, M.A., 2016. Biology and structure of leukocyte beta 2 integrins and their role in inflammation. F1000Res 5.

Baker, P.J., Boucher, D., Bierschenk, D., Tebartz, C., Whitney, P.G., D’Silva, D.B., Tanzer, M.C., Monteleone, M., Robertson, A.A., Cooper, M.A., Alvarez-Diaz, S., Herold, M.J., Bedoui, S., Schroder, K., Masters, S.L., 2015. NLRP3 inflammasome activation downstream of cytoplasmic LPS recognition by both caspase-4 and caspase-5. Eur J Immunol 45, 2918–2926.

Boltana, S., Roher, N., Goetz, F.W., Mackenzie, S.A., 2011. PAMPs, PRRs and the genomics of gram negative bacterial recognition in fish. Dev Comp Immunol 35, 1195–1203.

Fuhlbrigge, R.C., Chaplin, D.D., Kiely, J.M., Unanue, E.R., 1987. Regulation of interleukin 1 gene expression by adherence and lipopolysaccharide. J Immunol 138, 3799–3802.

Gahmberg, C.G., Valmu, L., Fagerholm, S., Kotovuori, P., Ihanus, E., Tian, L., Pessa-Morikawa, T., 1998. Leukocyte integrins and inflammation. Cell Mol Life Sci 54, 549–555.

Ghosn, E.E., Yang, Y., Tung, J., Herzenberg, L.A., Herzenberg, L.A., 2008. CDllb expression distinguishes sequential stages of peritoneal B-1 development. Proc Natl Acad Sci U S A 105, 5195–5200.

Goetz, F.W., Iliev, D.B., McCauley, L.A., Liarte, C.Q., Tort, L.B., Planas, J.V., Mackenzie, S., 2004. Analysis of genes isolated from lipopolysaccharide-stimulated rainbow trout (*Oncorhynchus mykiss*) macrophages. Mol Immunol 41, 1199–1210.

Harris, E.S., McIntyre, T.M., Prescott, S.M., Zimmerman, G.A., 2000. The leukocyte integrins. J Biol Chem 275, 23409–23412.

Hogg, N., Cabañas, C., Dransfield, I., 1993. Leukocyte integrin activation, in:*al*., P.E.L.e. (Ed.), Structure, Function, and Regulation of Molecules Involved in Leukocyte Adhesion. Springer-Verlag New York Inc..

Hughes, A.L., 2006. Evolutionary relationships of vertebrate NACHT domain-containing proteins. Immunogenetics 58, 785–791.

Humphries, J.D., Byron, A., Humphries, M.J., 2006. Integrin ligands at a glance. J Cell Sci 119, 3901–3903.

Iliev, D.B., Liarte, C.Q., MacKenzie, S., Goetz, F.W., 2005a. Activation of rainbow trout (*Oncorhynchus mykiss*) mononuclear phagocytes by different pathogen associated molecular pattern (PAMP) bearing agents. Mol Immunol 42, 1215–1223.

Iliev, D.B., Roach, J.C., Mackenzie, S., Planas, J.V., Goetz, F.W., 2005b. Endotoxin recognition: in fish or not in fish? FEBS Lett 579, 6519–6528.

Ingalls, R.R., Golenbock, D.T., 1995. CD11c/CD18, a transmembrane signaling receptor for lipopolysaccharide. J Exp Med 181, 1473–1479.

Johnson, M.S., Lu, N., Denessiouk, K., Heino, J., Gullberg, D., 2009. Integrins during evolution: evolutionary trees and model organisms. Biochim Biophys Acta 1788, 779–789.

Kettritz, R., Choi, M., Rolle, S., Wellner, M., Luft, F.C., 2004. Integrins and cytokines activate nuclear transcription factor-kappaB in human neutrophils. J Biol Chem 279, 2657–2665.

Latz, E., Xiao, T.S., Stutz, A., 2013. Activation and regulation of the inflammasomes. Nat Rev Immunol 13, 397–411.

Lüderitz, O., Galanos, C., Rietschel, E.T., 1982. Endotoxins of gram-negative bacteria. Pharmacol Ther 15, 383–402.

Miller, S.I., Ernst, R.K., Bader, M.W., 2005. LPS, TLR4 and infectious disease diversity. Nat Rev Microbiol 3, 36–46.

Nagai, Y., Akashi, S., Nagafuku, M., Ogata, M., Iwakura, Y., Akira, S., Kitamura, T., Kosugi, A., Kimoto, M., Miyake, K., 2002. Essential role of MD-2 in LPS responsiveness and TLR4 distribution. Nat Immunol 3, 667–672.

Petrilli, V., Dostert, C., Muruve, D.A., Tschopp, J., 2007. The inflammasome: a danger sensing complex triggering innate immunity. Curr Opin Immunol 19, 615–622.

Pfaffl, M.W., 2001. A new mathematical model for relative quantification in real-time RT-PCR. Nucleic Acids Res 29, e45.

Pfaffl, M.W., Horgan, G.W., Dempfle, L., 2002. Relative expression software tool (REST (c)) for group-wise comparison and statistical analysis of relative expression results in real-time PCR. Nucleic Acids Research 30.

Poltorak, A., He, X., Smirnova, I., Liu, M.Y., Van Huffel, C., Du, X., Birdwell, D., Alejos, E., Silva, M., Galanos, C., Freudenberg, M., Ricciardi-Castagnoli, P., Layton, B., Beutler, B., 1998. Defective LPS signaling in C3H/HeJ and C57BL/10ScCr mice: mutations in *Tlr4* gene. Science 282, 2085–2088.

Reis, M.I., do Vale, A., Pereira, P.J., Azevedo, J.E., Dos Santos, N.M., 2012. Caspase-1 and IL-1beta processing in a teleost fish. PLoS One 7, e50450.

Sakamaki, K., Satou, Y., 2009. Caspases: evolutionary aspects of their functions in vertebrates. J Fish Biol 74, 727–753.

Schmittgen, T.D., Livak, K.J., 2008. Analyzing real-time PCR data by the comparative CT method. Nature Protocols 3,1101–1108.

Schroder, K., Tschopp, J., 2010. The inflammasomes. Cell 140, 821–832.

Schutte, R.J., Parisi-Amon, A., Reichert, W.M., 2009. Cytokine profiling using monocytes/macrophages cultured on common biomaterials with a range of surface chemistries. J Biomed Mater Res A 88, 128–139.

Scott, M.J., Billiar, T.R., 2008. Beta2-integrin-induced p38 MAPK activation is a key mediator in the CD14/TLR4/MD2-dependent uptake of lipopolysaccharide by hepatocytes. J Biol Chem 283, 29433–29446.

Shi, J., Zhao, Y., Wang, Y., Gao, W., Ding, J., Li, P., Hu, L., Shao, F., 2014. Inflammatory caspases are innate immune receptors for intracellular LPS. Nature 514, 187–192.

Sollberger, G., Strittmatter, G.E., Garstkiewicz, M., Sand, J., Beer, H.D., 2014. Caspase-1: the inflammasome and beyond. Innate Immun 20,115–125.

Sunagawa, S., Coelho, L.P., Chaffron, S., Kultima, J.R., Labadie, K., Salazar, G., Djahanschiri, B., Zeller, G., Mende, D.R., Alberti, A., Cornejo-Castillo, F.M., Costea, P.I., Cruaud, C., d’Ovidio, F., Engelen, S., Ferrera, I., Gasol, J.M., Guidi, L., Hildebrand, F., Kokoszka, F., Lepoivre, C., Lima-Mendez, G., Poulain, J., Poulos, B.T., Royo-Llonch, M., Sarmento, H., Vieira-Silva, S., Dimier, C., Picheral, M., Searson, S., Kandels-Lewis, S., Tara Oceans, c., Bowler, C., de Vargas, C., Gorsky, G., Grimsley, N., Hingamp, P., Iudicone, D., Jaillon, O., Not, F., Ogata, H., Pesant, S., Speich, S., Stemmann, L·, Sullivan, M.B., Weissenbach, J., Wincker, P., Karsenti, E., Raes, J., Acinas, S.G., Bork, P., 2015. Ocean plankton. Structure and function of the global ocean microbiome. Science 348, 1261359.

Takle, H., Andersen, O., 2007. Caspases and apoptosis in fish. Journal of Fish Biology 71, 326–349.

Tumbol, R.A., Baiano, J.C.F., Barnes, A.C., 2009. Differing cell population structure reflects differing activity of Percoll-separated pronephros and peritoneal leucocytes from barramundi (*Lates calcarifer*). Aquaculture 292, 180–188.

Walzog, B., Weinmann, P., Jeblonski, F., Scharffetter-Kochanek, K., Bommert, K., Gaehtgens, P., 1999. A role for beta(2) integrins (CD11/CD18) in the regulation of cytokine gene expression of polymorphonuclear neutrophils during the inflammatory response. FASEB J 13, 1855–1865.

Weiss, A., Wiskocil, R.L., Stobo, J.D., 1984. The role of T3 surface molecules in the activation of human T cells: a two-stimulus requirement for IL 2 production reflects events occurring at a pre-translational level. J Immunol 133,123–128.

Wong, K.F., Luk, J.M., Cheng, R.H., Klickstein, L.B., Fan, S.T., 2007. Characterization of two novel LPS-binding sites in leukocyte integrin betaA domain. FASEB J 21, 3231–3239.

Wright, S.D., Jong, M.T., 1986. Adhesion-promoting receptors on human macrophages recognize *Escherichia coli* by binding to lipopolysaccharide. J Exp Med 164,1876–1888.

Wright, S.D., Ramos, R.A., Tobias, P.S., Ulevitch, R.J., Mathison, J.C., 1990. CD14, a receptor for complexes of lipopolysaccharide (LPS) and LPS binding protein. Science 249, 1431–1433.

Yi, E.C., Hackett, M., 2000. Rapid isolation method for lipopolysaccharide and lipid A from gram-negative bacteria. Analyst 125, 651–656.

Zoccola, E., Delamare-Deboutteville, J., Barnes, A.C., 2015. Identification of barramundi (*Lates calcarifer*) DC-SCRIPT, a specific molecular marker for dendritic cells in fish. PLoS One 10, e0132687.

Zoccola, E., Kellie, S., Barnes, A.C., 2017. Immune transcriptome reveals the mincle C-type lectin receptor acts as a partial replacement for TLR4 in lipopolysaccharide-mediated inflammatory response in barramundi (*Lates calcarifer*). Molecular Immunology 83, 33–45.

